# Reticulate evolution is favored in influenza niche switching

**DOI:** 10.1101/033514

**Authors:** Eric J. Ma, Nichola J. Hill, Justin Zabilansky, Kyle Yuan, Jonathan A. Runstadler

**Affiliations:** Department of Biological Engineering, Massachusetts Institute of Technology, 77 Massachusetts Ave, Cambridge, MA, 02139

**Author notes:** Corresponding Authors: Jonathan A. Runstadler,; Eric J. Ma.

**Keywords:** Ecology, Reticulate Evolution, Influenza

## Abstract

Reticulate evolution is thought to accelerate the process of evolution beyond simple genetic drift and selection, helping to rapidly generate novel hybrids with combinations of adaptive traits. However, the long-standing dogma that reticulate evolutionary processes are likewise advantageous for switching ecological niches, as in microbial pathogen host switch events, has not been explicitly tested. We use data from the influenza genome sequencing project and a phylogenetic heuristic approach to show that reassortment, a reticulate evolutionary mechanism, predominates over mutational drift in transmission between different host species. Moreover, as host evolutionary distance increases, reassortment is increasingly favored. We conclude that the greater the quantitative difference between ecological niches, the greater the importance of reticulate evolutionary processes in overcoming niche barriers.

**Significance Statement:** Are the processes that result in the exchange of genes between microbes quantitatively advantageous for those microbes when switching between ecological niches? To address this question, we consider the influenza A virus as a model microbe, with its ability to infect multiple host species (ecological niches) and undergo reassortment (exchange genes) with one another. Through our analysis of sequence data from the Influenza Research Database and the Barcode of Life Database, we find that the greater the quantitative difference between influenza hosts, the greater the proportion of reassortment events were found. More broadly, for microbes, we infer that reticulate evolutionary processes should be quantitatively favoured when switching between ecological niches.

## Manuscript

Reticulate evolutionary processes, such as horizontal gene transfer (HGT) and genomic reassortment, have been proposed as a major mechanism for microbial evolution (1), aiding in the diversification into new ecological niches (2). In contrast to clonal adaptation through genetic drift over time, reticulate evolutionary processes allow an organism to acquire independently evolved genetic material that can confer new fitness-enhancing traits. Examples include the acquisition of cell surface receptor adaptations (point mutations) in viruses (3), and antibiotic resistance (single genes) (4) and pathogenicity islands (or gene clusters) in bacteria (5). Host switching, defined as a pathogen moving from one host species into another, represents a fitness barrier to microbial pathogens. The acquisition of adaptations through reticulate processes either prior to or after transmission from one species to another may serve to aid successful pathogen host switches by improving fitness and the likelihood of continued transmission (6). In this sense, reticulate evolution may be viewed as an ecological strategy for switching between ecological niches (such as different host species), complementing but also standing in contrast to the clonal adaptation of a microbial pathogen by genetic drift under selection. In order to test this idea and its importance in host switch events (which are critical for (re-)emerging infectious disease), we provide a quantitative assessment of the relative importance of reticulate processes versus clonal adaptation in aiding the ecological niche switch of a viral pathogen.

Data yielded from influenza genome sequencing projects provides a unique opportunity for quantitatively testing this concept, and is suitable for the following reasons. Firstly, the influenza A virus (IAV) has a broad host tropism (7), and is capable of infecting organisms spanning millennia of divergence on the tree of life. With different host-specific restriction factors forming an adaptive barrier, each host species may then be viewed as a unique ecological niche for the virus (8). Secondly, IAV is capable of and frequently undergoes reassortment, which is a well documented reticulate evolutionary process (9-12). Reassortment has also been implicated as an adaptive evolutionary mechanism in host switching (13-14), though this is most prevalently observed for pandemic viruses of public health interest for which sequences are available (15). Finally, due to surveillance efforts over the past two decades, whole genome sequences have been intensively sampled over a long time frame, with corresponding host species metadata, available in an easily accessible and structured format (16). Because reassortant viruses are the product of two or more genetically distinct viruses co-infecting the same host, a more complex process than clonal transmission and adaptation, they are expected to occur less frequently. Hence, the global IAV dataset, which stretches over time and space with large sample numbers, provides the necessary scope to detect reassortant viruses at a scale required to quantitatively assess the relative importance of reticulate events in viral host switching.

In order to identify heterlologous reassortment events (between distinct influenza lineages) and the hosts species involved, we adapted a phylogenetic heuristic method (17), and mapped out a network of clonal and reassortment descent relationships from a global set of completely sequenced IAV (18,632 viral genomes) downloaded from the Influenza Research Database (16). Briefly, the core logic of the method is as such: for every isolate in the dataset, we look for genomic sources such that the sources found are of maximal similarity across all 8 genomic segments (Materials and Methods). Clonal descent involves tracing sources of whole genomes, while reassortment descent involves identifying source pairs, in which some segments of a sink virus’ genome comes from one source and a complementary set of segments comes from another source. Where either multiple sources or multiple source pairs correspond to the maximal similarity, all are kept as plausible sources, with appropriate weighting applied to avoid double-counting reassortment events (Materials & Methods). In the resulting network, nodes are individual viral isolates, and edges are the clonal or reassortment descent relationships.

In this network of viral isolates, clonal descent is mostly structured by host species, with known global patterns of human-to-human (H3N2 & H1N1, and rarer H5N1 & H7N9), chicken-to-chicken (H9N2, H7N9, H5N1) and swine-to-swine (H3N2, H1N1, H1N2) viral circulation captured in the network reconstruction (Figure S1). Edges in the network connected viral isolates with a median genetic similarity of 99.7%, indicating a high degree of genetic similarity captured in the network-based reconstruction (Figure S2). As expected, no clonal descent was identified between viruses of different subtypes. Moreover, the network recreates the phylogeny of known reassortant viruses, including the 2009 pandemic H1N1 and the recent 2013 H7N9 viruses, further validating the accuracy of our reconstruction (a browser-based d3.js visualization is available in Zenodo archive of the Github repository (Materials & Methods)). Small-world simulation studies validated our method as being accurate in detecting reassortment events (Figure S4), while a comparison of edges to a phylogenetic reconstruction on a subset of the data show that our method captures the shorter end of the distribution of patristic distances on a tree, indicating accurate approximation to phylogenetic reconstruction (Figure S3). Hence, our method is capable of detecting reassortment events, which are classically inferred by observing incongruences in phylogenetic tree clustering.

To test whether reassortment or clonal descent was an advantageous strategy when switching hosts, we computed the weighted proportion of reassortant edges (out of all edges) occurring between hosts of the same or different species. When host species were different, reassortant edges were over-represented at 19 percentage points above a null permutation model (permutation test described in Materials & Methods) (Figure 1a), and when host species were the same, reassortant edges were under-represented by 7 percentage points relative to our null model. Thus, reassortment is a strongly favored strategy when influenza crosses between different host species.

We further sought to explore whether the predominant use of reticulate evolutionary processes in host switch events were correlated with host phylogenetic relatedness and host ecology. To do this, we first computed the proportion of reassortment when switching between birds, non-human mammals, or humans, which are 3 divergent host groupings with distinct ecological behaviour. (For example, humans are the only known species to employ disease control measures, and affect the ecology of other species (birds and mammals through domestication) at scale.) We further sub-divided avian and mammalian categories into wild and domestic, to assess the impact of anthropological activity on the relative importance of reassortment in host switch interfaces (see Materials and Methods for how AIV was classified as domestic or wild). To ensure that the dataset was sufficient in scope to detect reassortant viruses, we only considered host group transitions with at least 1000 descent events (both clonal and reassortant), or at least 10 reassortment events (dashed yellow and green lines respectively in Figures 1b & c). Nonetheless, all data are displayed for completeness.

Here, reassortment is over-represented relative to the null when host groups are different. Only two exceptions occur. The first is between wild birds, where reassortment is over-represented but host groups are not different. In this case, the “wild bird” label encompasses a wide range of host species, and as the natural reservoir for many diverse influenza viral subtypes, we expect to detect reassortment events more frequently between diverse species that may be distantly evolutionarily related.

The second is the human-domestic mammal interface, where reassortment is not over-represented even though the host groups are different. In the case of human to domestic mammal host switches (reverse zoonosis), these are mostly well-documented reverse zoonotic events between human and swine hosts (18)), where shared cellular receptors for viral entry (19) facilitates zoonotic and reverse zoonotic transmission. This may be a case of host convergent evolution inadvertently lowering the adaptive barrier to host switching. Under representation of reassortment at human-to-human transitions is expected because of the limited number of viral subtypes circulating in human populations that undergo serial selective sweeps, resulting in high sequence similarity within the viral pool (20), which likely obscures the distinction between reassortment and clonal descent. However, we also expect antibody-mediated immunity, whether from vaccination or prior exposure, to further limit the frequency of co-infection and likelihood of reassortment events happening amongst humans. Thus, despite the exceptions that may be explained by our current best knowledge of influenza biology (e.g. human to swine transmissions), reassortment is strongly favored over clonal evolution when crossing between evolutionarily distant hosts.

To further explore the relationship between host evolutionary divergence and the predominance of reassortment in transmission events between species, we compared a common phylogenetic measure of species divergence, the cytochrome oxidase I (COI) gene, to the use of reassortment in host switch events. A subset of viral hosts, encompassing a variety of bird and mammal species, have had their cytochrome oxidase I (COI) gene sequenced as part of the barcode of life project (21). For the subset of edges in the network for which both the source and sink hosts have a COI gene sequence that fulfilled our criteria for consideration (as described above), we computed the percentage evolutionary distance between the two hosts (Materials and Methods). Applying a similar permutation test and assessment criteria as described for host groups above, we found a trend of increasing over-representation at higher evolutionary distances (Figure 1c). Thus, as host evolutionary distance, or more broadly, as quantitative niche dissimilarity increases, reticulate evolution becomes increasingly favored for influenza virus niche switch events.

We have quantitatively defined the importance of reticulate evolutionary events in switching ecological niches, using an infectious disease data set with characteristics that are particularly well suited for answering this question. Beyond the viral world, recent reviews have asserted the importance of reticulate evolutionary events as a driver of speciation and niche diversification (22-23), and recent studies have illustrated heightened fitness effects in hybrid populations (24-25). However, none have quantitatively tested the importance of reticulate evolutionary strategies in enabling ecological niche switches at a global scale, especially in comparison to clonal adaptation under drift and selection (a task feasible only in fast evolving organisms). Additionally, no studies to date have examined reticulate evolutionary processes in the context of quantified niche differences, as we have done here by measuring reassortment in the context of host evolutionary distance. Our study provides strong quantitative evidence supporting the hypothesis that reticulate evolutionary processes are advantageous relative to adaptation by drift for pathogen transfer between host species, and therefore more broadly, ecological niche switching.

We note four limitations to this study. Firstly, we recognize that in this study, we have considered only a single pathogen for which abundant genomic data are available, and whose genomic and host tropic characteristics are suitable for this analysis. To specifically answer whether reticulate processes are favored over clonal transmission for other organisms, using these methods, depends on being able to acquire genome sequences with matched ecological niche metadata.

Secondly, we also note that the global influenza dataset will have unavoidable sampling biases. For example, human isolates predominate in the dataset, and consequently the human-associated subtypes H3N2 and H1N1 also dominate the dataset. Sequences from viral outbreaks will also be over-represented relative to isolates collected through routine surveillance sampling, and will unavoidably lead to a heightened detection of clonal descent in a single host species. In order to deal with this sampling bias, our permutation tests (for the host species and group labels) involve class labels of equal sizes. This allows us to calculate an expected distribution of proportions under ideal assumptions of equal sampling, which in turn forms the baseline for our conclusions.

Thirdly, our choice to use “host species” as the defined and quantified ecological niche is, in part, borne out of data availability. We naturally expect exceptions to occur if differences between species do not constitute a *major* barrier, or if barriers are defined by other characteristics of the host. Mallards are one example of such an exception (Figure S7). Amongst mallards, pre-existing immunity (perhaps quantifiable by antibody landscapes (26)) and high subtype diversity may be a strong driving forces for reassortment (27). We note that the necessary data do not currently exist to quantify barriers for other levels of defining and quantifying ecological niches, such as individuals or populations, at a global scale.

Finally, we do not specifically identify whether reassortment occurs prior to or after host switching, but only identify host transitions across which reassortment is implicated. A reassortment event may occur within a host species, during transfer between two host species, or after the transfer; reassortment’s association with host switching will depend on when the reassortant virus is detected, and consequent clonal expansion of the reassortant strain will be identified as “clonally descended”. Our method does not identify when the reassortment event happens, and this is both a limitation of our method and of IAV surveillance being less dense than necessary to distinguish between these two scenarios. Without better prior knowledge on whether reassortment happens prior to or after host switching, our method assumes that the detected reassortment events are the best possible representation of ground truth. It is with this limitation in mind that we identify associations of reassortment events with host switches, or more broadly across ecological niches. Whether reticulate evolution is causal for ecological niche switching will require further study.

In summary, using data available from a model zoonotic viral pathogen, we have shown that reticulate evolutionary processes are important in enabling pathogen host switches. For the influenza virus, reticulate evolution predominates when crossing between hosts. More broadly, the greater the quantitative difference between ecological niches, the greater the importance of reticulate evolutionary processes in enabling niche switches. While the quantitative importance of reticulate evolution may differ for different organisms evolving in different niches, we expect that further sequencing efforts from across broad domains of microbial life, and a further characterization and definition of their ecological niches, will elucidate whether this principle holds more broadly. Beyond its relevance to evolutionary ecology, reticulate evolution also has public health consequences. Reassortant influenza viruses have been implicated in all past human pandemic strains for which we have sequence data (28-31), and the ancestry of HIV-1 involved a hybrid SIV (32). Hence, knowing how reticulate events shape disease emergence may help the ecology and evolution of infectious disease become a more predictive science, leading to insight important to disease prevention and mitigation (33).

## Acknowledgments

The authors would like to thank Dr. Nan Li for technical assistance on graph computation during the earlier stages of this work. We would like to acknowledge funding from the NIH/NIAID CEIRS Program (Contract no.: HHSN272014000008C) and the MIT Department of Biological Engineering. The authors would also like to thank William R. Hesse of the MIT BE Communications Lab for assistance in reviewing the manuscript.

## Materials and Methods

**Sequence data source and preprocessing**. Sequence and metadata was downloaded from the Influenza Research Database on 2 September 2015. Search parameters on the IRD were as such: Segment/Nucleotide data, Virus Type A, All Segments, All Hosts, All Geographic Groupings, Complete Genomes Only, Include pH1N1 sequences, Date Range 1980-Present. Advanced options included: Laboratory Strains excluded. Segment FASTA files were downloaded, with a Custom Format where only Accession Numbers were included.

**Source code and Data**. Digital Object Identifiers (DOIs) hosted on zenodo.org can be found for the following source code.

- Source code for graph construction: 10.5281/zenodo.33421.
- Source code for analysis, derivative data (network node and edge lists, computed threshold values) and figure construction as part of a series of Jupyter notebooks: <u>10.5281/zenodo.47267</u>.
- Source code for the patristic distance tests: 10.5281/zenodo.33425.
- Source code and Jupyter notebooks for simulation studies: 10.5281/zenodo.33427

**Reassortant virus identification**. We adapted the SeqTrack algorithm (17) to perform graph construction. Sequences were aligned using Clustal Omega 1.2.1 (34), and the resultant distance matrix was converted into a similarity matrix by taking 1 – *distance*. Affinity Propagation (35) clustering was done on each segment’s similarity matrix to determine a threshold cut-off similarity value, defined as the *minimum (across all clusters for that segment) of minimum incluster pairwise identities*, below which we deemed it implausible for an evolutionary descent (clonal or reassortment) to have occurred (Figure S6). Because the Affinity Propagation algorithm does not scale well with sample size, we treated the threshold computation as an estimation problem, and the final threshold was computed as the median threshold of 50 random sub-samples of 500 isolates.

We then thresholded each segment’s similarity matrix based on its segment’s threshold value, summed all 8 thresholded similarity matrices, and then for each isolate, we identified the most similar isolate that occurred prior to it in time. This yielded the initial “full complement” graph without reassortant viruses. Each edge in this graph has an attached PWI, which is the sum of PWIs across all 8 segments. Within this graph, there are isolates for which no “full complement” of segments could be identified, which are candidate reassortant viruses. Additionally, amongst the isolates for which a full complement of segments could be found from another source, we identified those whose in-edges were weighted at the bottom 10% of all edges present in the graph, which we also identified as candidate reassortant viruses (1357 out of 1368 of such viruses were eventually identified as reassortant; the other 11 were considered to be clonally descended). For these viruses, we performed source pair searches, where we identified sources for a part of the genome from one virus and sources for the complementary part of the genome from another virus. If the summed PWI across the segments for the two viruses was greater than the single-source search, we accepted the source pair as the candidate reassortant.

**Edge Weighting and Proportion Reassortment Calculations**. The proportion of reassortment events was calculated by first weighting each incoming edge to every virus. The weighting procedure is described here: If the virus is detected to be plausibly clonally descended from n other viruses, as determined by maximal similarity, it is given a weight of 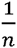. If the virus is detected to be a reassortant, then edges are weighted by the the fraction of times that it is involved in a max similarity source pair. (Figure S5) For example, if a given node A has a plausible source of segments in B, C, D, with (B and C) and (B and D) being plausible sources, then the edge B-A would be given a weight of 0.5, and the edge C-A and D-A would be given a weight of 0.25 each. (Figure S5) The proportion of reassortment edges was then calculated by taking the sum of weights across all reassortant edges (for a particular transition, e.g. between or within host species), divided by the sum of weights across all reassortant and clonal edges (for the same particular transition). With this weighting scheme, multiple plausible sources that lead to the same virus are not double-counted.

**Permutation Tests**. Null models were constructed by permuting node labels; equal class size permutation was performed for host species (for Figure 1a) and host group (for Figure 1b). For example, if there were 9 ‘human’, 4 ‘swine’ and 2 ‘chicken’ nodes (15 total), labels would be randomly shuffled amongst the nodes such that each label were equally represented (5 each).

**Host Group Labelling**. Host species were manually classified into the “human”, “domestic animal” and “wild animal” groups, based on the country of isolation. For example, “ducks” would be considered a “wild animal” in North America, while it would be considered a “domestic animal” in East Asian countries. Ambiguous host species, while remaining in the dataset, were excluded from the analysis.

**Host Evolutionary Distance**. Host species’ scientific names were sourced from the Tree of Life database (www.tolweb.org). Only host species with unambiguous scientific names recorded were considered. Cytochrome oxidase I genes were sourced from the Barcode of Life Database (www.boldsystems.org) on 31 October 2015. Sequences had to be at least 600 n.t. long to be considered, and only positions with fewer than 3 gap characters were concatenated into the final trimmed alignment. Evolutionary distance was computed from the trimmed alignment as the proportion of mismatched nucleotides. Further details are available in the Jupyter notebooks.

**Simulation Studies**. In our simulations, we sought to model the process of obtaining sequences as simply as possible. Therefore, we used a kinetic Monte Carlo-style process to simulate the generation of new viral sequences under the processes of replication, mutation and reassortment. Briefly, we simulated simple two-segment model viruses, with each of the two segments having a different substitution rate. Each simulation run was initialized with anywhere between 1 and 5 viruses. At each time step, one virus was chosen at random to replicate (with 0.75 probability), or reassort with another virus (with 0.25 probability). Simulations were run for 50 time steps. Regardless of replication or reassortment, the progeny virus is subjected to mutations, with the number of mutations in each segment being drawn from a Binomial distribution with probability equal to the segment’s substitution rate, and the exact positions drawn uniformly across the segment. Specific implementation is viewable in the IPython HTML notebooks and Python class definitions available on Github, available at the URL provided in Code and Data Deposition. This simulation process thus gives rise to a fully known ground-truth graph, which all reconstructions can be compared against.

The number of unique starting genotypes and total number of viral isolates being considered was much smaller than the real-world data. Therefore, our graph reconstruction procedure captured the essential parts of the method employed in the global analysis, but differed in the details. Here, “full complements” involve only two segments. We did not perform affinity propagation clustering as we started with completely randomly generated sequences of equal length. Our “null model” graph is where source isolates are chosen uniformly at random from the set of nodes occurring prior to the sink isolates.

In order to assess the accuracy of our reconstruction, we defined the path accuracy, and reassortant path identification accuracy metrics. Edge accuracy, which is not used for evaluation here, is whether a particular reconstruction transmission between two isolates exists in the simulation. Path accuracy is a generalization of edge accuracy, where a path existing between the source and sink nodes (without considering the direction of edges) in the reconstruction is sufficient for being considered accurate. Reassortant path identification accuracy measures how accurately we identified the reassortant paths, analogous to the regular path accuracy.

**Phylogenetic Reconstruction and Patristic Distance Comparison**. Phylogenetic reconstruction was done for a subset of H3N8 viruses isolated from Minto Flats, AK, between 2009 and 2010 as part of a separate study. Briefly, each segment of the viral genomes were individually aligned using Clustal Omega (34), and their genealogies reconstructed using BEAST 1.8.0 (36). A minimum of 3 MCMC runs that converged on a single optimal tree were chosen to compute the maximum clade credibility (MCC) tree. Burn-in ranged from 10 to 39 million steps out of 40, with median 24 million steps. Patristic distances were calculated using the DendroPy package (37). In the graph reconstruction on the Minto Flats study, we extracted the edges and nodes involving only the H3N8 isolates, and computed the tree patristic distances between isolate pairs linked by an edge in the graph.

## Figure Legends

Figure 1. **Reassortment is over-represented relative to clonal descent in transmission across host barriers**. Proportion of reassortment events when crossing between (a) different or same hosts, (b) different host groups, and (c) hosts of differing evolutionary distance as measured by divergence in the cytochrome oxidase I (COI) gene. Reassortment is over-represented relative to clonal descent in transmission across host barriers. (b) D: Domestic animal, H: Human, W: Wild, B: Bird, M: Mammal. Donor host is labeled first. Bolded x-axis tick labels indicate data for which the weighted sum of all edges exceeded 1000, or the weighted sum of reassortant edges exceeded 10. (c) Pairwise distances between host’s cytochrome I oxidase genes are binned in increments of 5%, or 0.05 fractional distance. (a, b, c) Vertical error bars on the null permutation model represent 3 standard deviations from the mean from 100 simulations (a, b), or 95% density intervals from 500 simulations (c). (b, c) Translucent dots indicate the weighted sum of all (clonal and reassortment) descent (yellow) and reassortment (green) events detected in the network under each host group transition. Horizontal yellow and green lines indicates threshold of values of 1000 and 10 respectively.

